# This microtubule does not exist: Super-resolution microscopy image generation by a diffusion model

**DOI:** 10.1101/2023.07.06.548004

**Authors:** Alon Saguy, Tav Nahimov, Maia Lehrman, Estibaliz Gómez-de-Mariscal, Iván Hidalgo-Cenalmor, Onit Alalouf, Ricardo Henriques, Yoav Shechtman

## Abstract

Generative models, such as diffusion models, have made significant advancements in recent years, enabling the synthesis of high-quality realistic data across various domains. Here, we explore the adaptation and training of a diffusion model on super-resolution microscopy images from publicly available databases. We show that the generated images resemble experimental images, and that the generation process does not memorize existing images from the training set. Additionally, we compare the performance of a deep learning-based deconvolution method trained using our generated high-resolution data versus training using high-resolution data acquired by mathematical modeling of the sample. We obtain superior reconstruction quality in terms of spatial resolution using a small real training dataset, showing the potential of accurate virtual image generation to overcome the limitations of collecting and annotating image data for training. Finally, we make our pipeline publicly available, runnable online, and user-friendly to enable researchers to generate their own synthetic microscopy data. This work demonstrates the potential contribution of generative diffusion models for microscopy tasks and paves the way for their future application in this field.

## Introduction

Deep learning algorithms have been extensively used in the past decade to solve various microscopy challenges^1–7^. These algorithms outperform traditional computer vision methods in terms of reconstruction quality, analysis time, and classification, among many others. However, deep learning solutions are hungry for data. To train a model, one should typically acquire and annotate hundreds or even thousands of images manually, a highly time and resource consuming process. An alternative approach is to produce synthetic data based on mathematical models that describe the structure of the biological specimen^1,3,7–9^. Yet, tuning the data generation parameters is a cumbersome process that leads to non-realistic features in the synthetic images due to parameter estimation errors and model inaccuracies, which is critical to train highly generalizable and accurate models.

Recently, the field of generative models has seen a significant surge in terms of both development and application^10–13^. Generative models have moved far beyond their initial application in producing artificial images and are now being used to create synthetic datasets that can effectively mimic real-world data in diverse domains^14^. Two major contributors to this advancement have been Denoising Diffusion Probabilistic Models (DDPM)^10^ and Denoising Diffusion Implicit Models (DDIM)^13^. DDPM and DDIM offer a dynamic approach for the generation of synthetic data, relying on stochastic processes to create totally new images that still capture the inherent image statistics present in the training dataset.

The capacity of diffusion models to accurately create realistic visual data is profoundly impacting many computer vision applications^15^, including microscopic imaging, where overcoming the existing challenges to gather high-quality large training datasets is invaluable. Indeed, several studies already incorporate diffusion models to microscopy to reconstruct 3D biomolecule structures in Cryo-EM images^16^, predict 3D cellular structures out of 2D images^17^, or drug molecule design^18^, among others.

Here, we propose the application of generative diffusion models in the field of super-resolution microscopy. First, we show the ability of diffusion models to generate realistic, high-quality, super-resolution microscopy images of microtubules and mitochondria. Then, we assess the capacity of the models to learn the intricate nature of the data domain by validating that the network does not memorize images from the training data. Next, we utilize the generated dataset to train a single-image super-resolution deep learning model and show superior reconstruction quality compared to the same model trained on model-based simulated data. The diffusion model approach proposed here is publicly available^19^ on the ZeroCostDL4Mic platform^20^, enabling non-expert researchers to benefit from it.

## Results

We base our work on a previously reported^21^ diffusion model which we adapt to super-resolution microscopy. We trained two diffusion models on different biological samples, microtubules and mitochondria, sourced from a publicly available database (ShareLoc.xyz^22–24^). We split our data into 60% and 40% for training and validation of the performance, obtaining a 7:5 training:validation image ratio for the microtubule data and a 3:2 training:validation image ratio for the mitochondria data. Furthermore, we split each image into patches of 256×256 pixels and transformed them using random horizontal flips and rotations of 90, 180, and 270 degrees to augment the training data. The augmentation step yielded a total of 2000 training patches for the microtubule data and 800 training patches for the mitochondria data. Training details are further specified in the Methods section.

The images generated by our DDPM qualitatively resemble the training data used for the training, as can be clearly seen in the examples in Figure 1. To validate that our model does not memorize images, namely, copy existing images from the training set and generate them as network outputs, we calculated the normalized cross-correlation between every generated image (a total of 50 images), including rotated and flipped versions of the images, and the augmented patches used for training. The maximal normalized cross-correlation, calculated between all generated images and the training data, was 0.345 (0.682) for the synthetic microtubule (mitochondria) images. The mean normalized cross-correlation value was 0.336 (0.631) for the microtubules (mitochondria) images. For comparison, repeating this process of normalized cross-correlation calculation between experimental images, taken from different datasets (a total of 10 images), and all other training data, yielded a mean value of 0.372 (0.414) and max value of 0.483 (0.510) for the microtubule (mitochondria) data. Then, we overlaid the training images that obtained the highest cross-correlation score with the generated images to verify that our synthetic images are sufficiently new and different from the training images (Figure 2).

**Figure 1:**
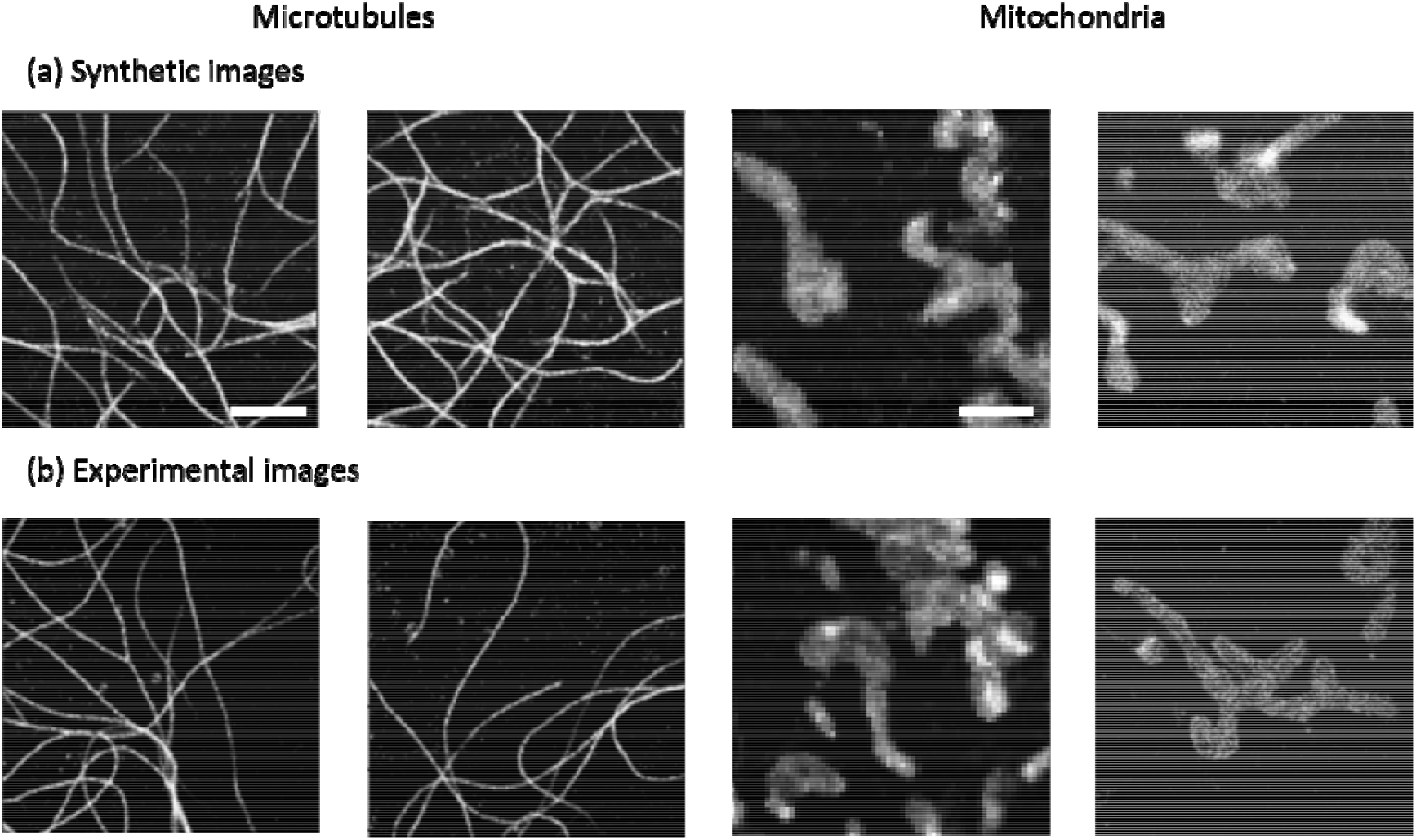
Qualitative comparison of experimental microscopy data versus data generated using our generative diffusion model. (a) Example synthetic images of microtubules (alpha-tubulin – Alexa647) and mitochondria (TOM 22 – Alexa647) generated by our diffusion model. (b) Example experimental super-resolution images, used as training data. Scale bars = 2.5 .

**Figure 2:**
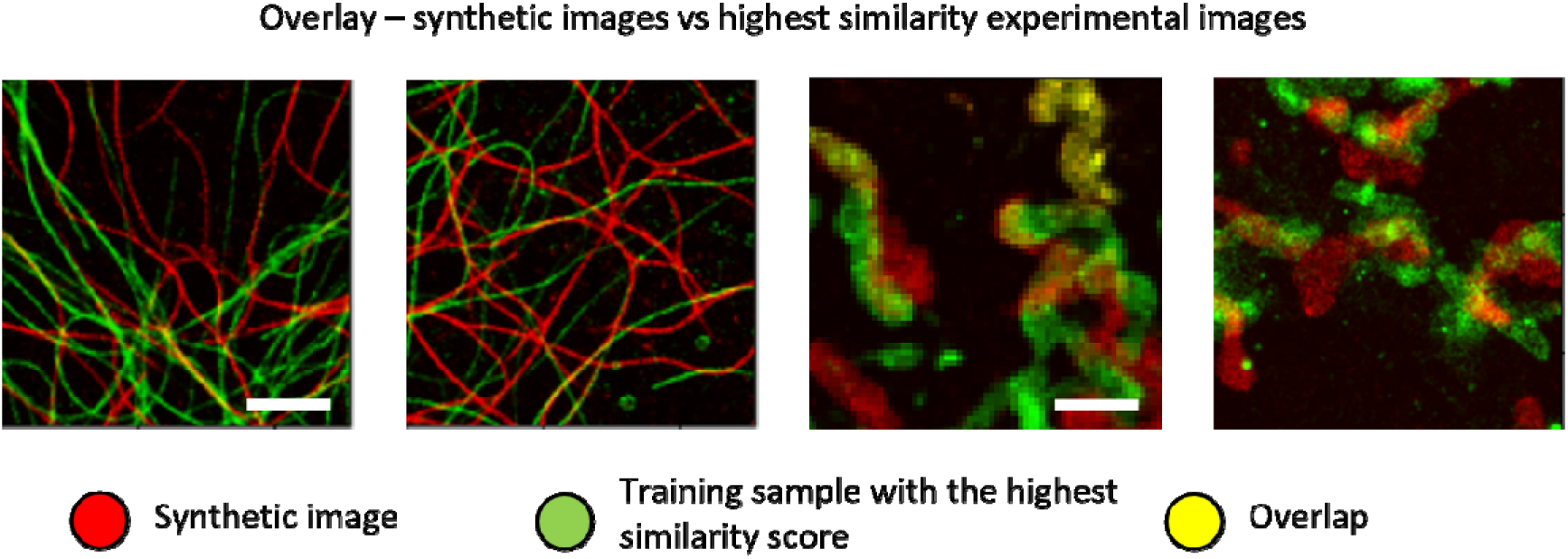
Diffusion models do not memorize training images. Overlay between each reconstructed image and the training image with highest resemblance (maximal cross-correlation score). Red marks generated data, green marks the closest training sample, and yellow marks overlap between the two images. Scale bars = 2.5 .

Notably, the cross-correlation values are similar for experimental microtubule images from another dataset (imaged in similar conditions) and the microtubule images that were generated by our diffusion model, showing the expected variability between different and independent datasets. In the case of mitochondria images, the cross-correlation values were slightly higher than those obtained when comparing with images from a different experimental dataset (see in-depth analysis in the discussion section).

Next, we tested the applicability of our generated data to improve deep learning-based methods. We used our generated data to train an application of Content-Aware Restoration (CARE)^1^, specifically, a deconvolution method aiming to transform a low-resolution image to a high-resolution image based on prior knowledge of image-statistics. Notably, obtaining single-image-based super-resolution algorithmically is yet an unsolved problem in microscopy, with results strongly dependent on the prior information provided, and is no match to physics based super-resolution microscopy methods (SMLM, STED, SIM, etc.^25–28^). Nevertheless, we use this task to demonstrate the potential of diffusion model-based data generation in virtual super-resolution microscopy imaging. We trained two CARE models for each biological sample using 1) synthetic images generated by a mathematical model and 2) images generated by our diffusion model. During the training stage, we simulated synthetic high-resolution images either by our model or a mathematical model; next, we obtained low-resolution images by forward passing the high-resolution images through a model of our optical system (see methods section for more details). Ultimately, we used these low-resolution – high-resolution pairs to train CARE.

Visually, the CARE model trained on data generated by the diffusion model yielded a better reconstruction in comparison to the traditionally trained network (Figures 3, 4). Moreover, we have analyzed the spatial resolution we obtained in both reconstructions using the Fourier Ring Correlation (FRC) plug-in for ImageJ^29^. In brief, FRC is a similarity measure that seeks the maximal spatial frequency in which the reconstructed and ground truth images are similar up to a predefined threshold. The similarity is quantified by the normalized cross-correlation between the Fourier transforms of both images inside a torus with increasing radius. A high cross-correlation value within the torus indicates high similarity between the images, in the corresponding spatial frequency band.

**Figure 3:**
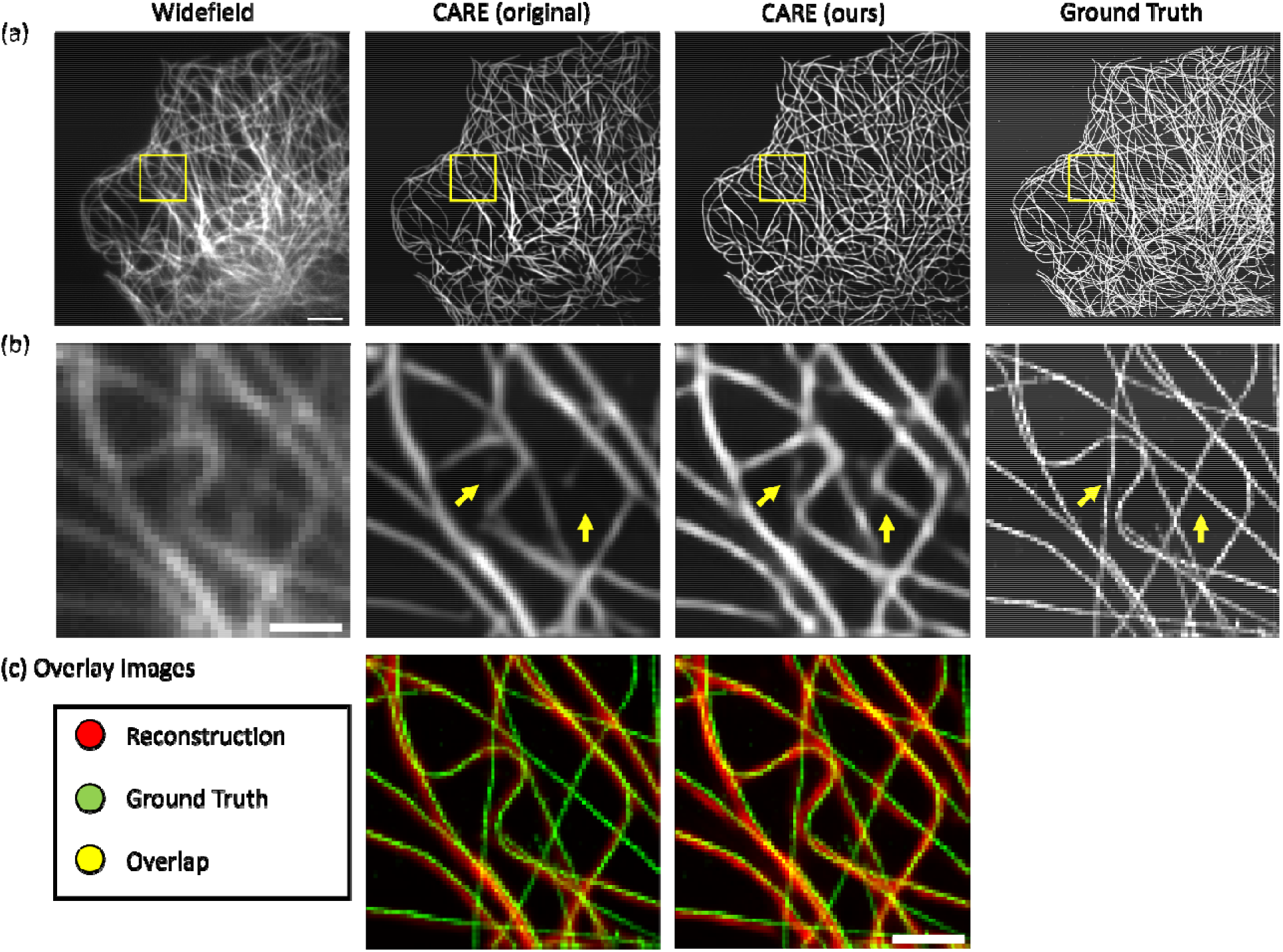
Performance of CARE trained on synthetic microtubule images generated by a mathematical model vs. training on microtubules generated by our diffusion model. (a) Left to right: widefield image, CARE reconstruction when trained on mathematical simulations, CARE reconstruction when trained on our synthetic data, and ground truth. Scale bar = 5 . (b) Regions of interest (marked by yellow squares in (a)), yellow arrows mark areas in which CARE trained on our data outperformed the previous method. (c) Left: overlay between CARE trained on mathematical simulations (red) and the ground truth (green). Right: overlay between CARE trained on our diffusion model-based synthetic data (red) and the ground truth (green). Scale bar = 1 .

**Figure 4:**
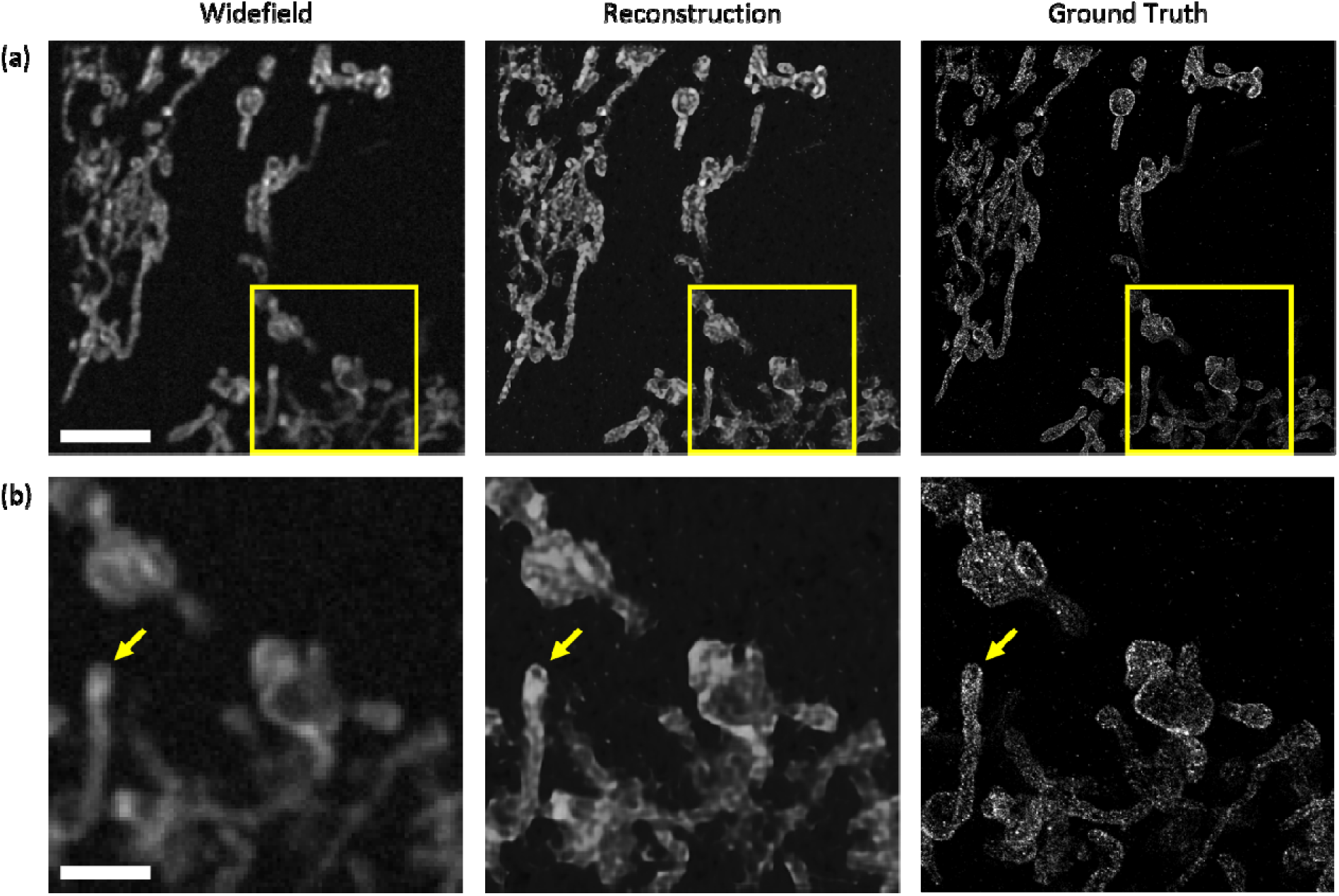
Performance of CARE trained on mitochondria generated by our diffusion model. (a) Left to right: widefield image, CARE reconstruction when trained on our synthetic data, and ground truth. Scale bar = 5 . (b) Region of interest; yellow arrow marks a subtle feature not visible in widefield imaging, which is made visible in our reconstruction. Scale bar = 2 .

The mean spatial resolution of the reconstructed images, as quantified by FRC, using a 1/7 threshold^29^ when training on microtubule images generated by our diffusion model was 100 nm, while the mean spatial resolution obtained when training on synthetic microtubules generated via a mathematical model was 140 nm.

Notably, microtubule images can be simulated with relatively high fidelity by a variety of well-established mathematical models^30^. However, for an arbitrary type of biological specimen, it is not easy to obtain a simple mathematical model describing its shape and characteristics. Therefore, the most remarkable feature of diffusion model-based data generation is the ability to generate synthetic data from non-mathematically defined biological specimen. Additionally, diffusion models might also contribute to the understanding of biological structures in a data driven manner by interpretation of the generated image statistics.

We validate this claim by training our diffusion model on publicly available super-resolution images of mitochondria^23^ (Figure 4). The spatial resolution obtained by CARE trained on our generated mitochondria was 110 nm. Unlike for microtubules, there is no available mathematical model to generate mitochondria images. Therefore, training CARE traditionally would require obtaining enough extended super-resolution images.

## Discussion

In this work, we demonstrate the potential of diffusion models to generate large super-resolution microscopy datasets by relying on a relatively small number of super-resolution images. Given only 7 microtubule images we managed to generate realistic microtubule images that looked different from the original training data, while resembling similar experimental data distribution. Notably, when training on yet a smaller dataset, i.e., 3 mitochondria images, some parts of the image were memorized, yielding slightly higher cross-correlation values when using the generated images than the values obtained when using mitochondria images from a different dataset. This observation is in par with existing work in this field^31,32^, implying that larger training sets prevent memorization and increase the uniqueness of the generated data. Therefore, we suggest using quantitative sanity checks (such as the cross-correlation metric) on the generated data as a tool to evaluate whether enough images were used to train the diffusion model. Additionally, when choosing the number of images for training, one should also take into consideration the data complexity and the size of the field-of-view of each image.

Creating synthetic images of biological data that are highly realistic and representative of the original data has important implications. For example, diffusion models enable efficient generation of super-resolution datasets that could be transformed to low-resolution observations by forward passing through an optical model of the imaging system; then, one may perform supervised model training without the need for extensive experimental data acquisition, often a limiting factor due to the impractical duration of the acquisition process. The contribution of our method is particularly relevant for the general case where no simple mathematical model is available for synthetic image generation.

Moreover, the ability to learn the complexity of biological structures and reproduce them to create realistic simulated images is key to identifying and interpreting biological phenomena and reinforcing data-driven discoverability. Our experiments have shown that diffusion models can be of great value in this area; these models have the ability to produce synthetic images that are of high quality and closely resemble real microscopy data, even when dealing with complex structures such as microtubule networks. To obtain accurate and highly generalizable models, it is essential to train deep learning models with the most realistic and extensive data possible, covering most of the natural experimental domain. These highly effective models, also known as foundation models, require massive amounts of image data to be trained effectively, which could be partially alleviated by smart and accurate data generation.

While the task chosen in this work to demonstrate the potential of the approach is single-image super-resolution, the applicability of diffusion model-based image generation for microscopy is naturally much broader. Numerous potential applications exist, including denoising, multi-image super-resolution, cross-modality imaging, live-cell dynamic imaging, and more. On the other side, quantitative evaluation of biological image data generation in the lack of annotated images is still an open question in the field that requires further work and consensus.

We share an easy-to-use notebook via the ZeroCostDL4Mic^20^ platform to enable researchers to replicate our pipeline and harness diffusion model capabilities. We also distribute the pretrained models that allow the generation of data similar to the data presented in this work. Of note, training diffusion models is time consuming due to the large number of stochastic operations involved in the learning process.

In light of the encouraging results obtained from this study, future research should continue to focus on further optimizing and evaluating diffusion models for generating more types of synthetic microscopy data and on finding the applications where these capabilities are most impactful. Furthermore, due to the capacity of diffusion models to create virtual representations of nanoscale cellular structure, they can potentially predict prospective multi-structural spatial relationships that will guide observations and discovery in the field of microscopy. The emergence of generative models for microscopy represents an exciting phase for bio-medical research and holds promising potential for advancements in the near future.

## Methods

### Optical model for low-resolution image generation

To train CARE on low-resolution – high-resolution image pairs, we used high-resolution data and passed it through a model of our optical system to obtain low-resolution images. In this work, we use a simple model to simulate a 2D low-resolution image based on a 2D high-resolution image. Let the imaged structure be depicted by *S(x*,*y)* and let *H(x*,*y)*, the point spread function (PSF) of the optical system, be modeled as a 2D Gaussian:

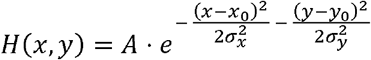

Where *A* is the amplitude of the PSF, *x*_0_, *y*_0_ are the position of the emitter, and *σ*_*x*_ *= σ*_*y*_ *= σ* represents the PSF width.

The low-resolution image formed at the camera is described by the convolution of the imaged structure with the system’s PSF equation:

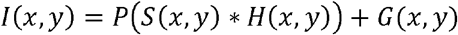

Where * indicates a convolution operator, *P(x, y)* indicates a Poisson distribution of the emitted number of photons, and *G(x, y)* indicates a Gaussian noise simulating the camera read noise.

### Diffusion model architecture and training details

We have adopted the network architecture presented by Nichol, et al^21^ We used a single residual network (ResNet) block and we changed the input and output layers of the model to fit monochromatic data. To decrease the network size, we also changed the channel multiplication between different layers of the ResNet, namely, instead of (1, 1, 2, 2, 4, 4) multiplication we used (1, 1, 2, 2, 2, 2) multiplication, where the initial channel number is 64. Additionally, we changed the number of diffusion steps to 2000, set the batch size to 10, the learning to 1*e*^-5^, and employed a cosine noise schedule. To train the network, we used 7 super-resolution localization lists of microtubule experiments and 3 of mitochondria experiments, all publicly available (ShareLoc^22^); then, we generated from each localization list a super-resolved image scaled by a factor of 4 in comparison to the diffraction limited data, yielding pixel sizes of 27 nm and 32 nm for the microtubule and mitochondria images, respectively.

Next, we split the input images to multiple overlapping patches of size 256*x*256 *pixels*^2^ and augmented the patches by flipping and rotating the images horizontally and vertically. The total number of training patches we used is 2000 and 800 for the microtubule and mitochondria networks respectively. Finally, we trained the generative diffusion model over 80,000 steps for 8 hours on a single NVIDIA 32GB Titan RTX GPU. Ultimately, generation of a single super-resolution image depends on the image size, e.g. 30 seconds for images of size 256*x*256 pixels^2^.

### CARE training details

We obtained super-resolution training data based on: 1) the mathematically simulated data presented in CARE paper; 2) the data generated by our trained diffusion model. To generate the low-resolution data needed for training CARE network, we followed a similar scheme as described in the CARE paper by convolving the super-resolution data with a gaussian microscope PSF model and adding Perlin noise, shot noise and gaussian noise. Importantly, we made sure that images generated by the two methods described above shared properties such as signal-to-noise ratio, sample size, etc. Finally, we trained the CARE network on 5000 synthetic low-resolution-high-resolution image pairs for the microtubule reconstruction and 2000 for the mitochondria reconstruction. To maintain a fair comparison between CARE trained on our data vs CARE trained on the mathematically generated microtubules, we used the same training set size in both cases.

